# Tim-3 co-stimulation promotes short-term effector T cells, restricts memory precursors and is dispensable for T cell exhaustion

**DOI:** 10.1101/179002

**Authors:** Lyndsay Avery, Andrea L. Szymczak-Workman, Lawrence P. Kane

## Abstract

Tim-3 is highly expressed on a subset of T cells during T cell exhaustion, in settings of chronic viral infection and tumors (1, 2). Using LCMV Clone-13, a model for chronic infection, we have found that Tim-3 is neither necessary nor sufficient for the development of T cell exhaustion. Nonetheless, expression of Tim-3 was sufficient to drive resistance to PD-L1 blockade therapy during chronic infection. Strikingly, expression of Tim-3 promoted development of short-term effector T cells, at the expense of memory precursor development, after acute LCMV infection. These effects were accompanied by increased Akt/mTOR signaling in T cells expressing endogenous or ectopic Tim-3. Conversely, Akt/mTOR signaling was reduced in effector T cells from Tim-3 deficient mice. Thus, Tim-3 is essential for optimal effector T cell responses, but may also contribute to exhaustion, by restricting development of long-lived memory T cells, including PD-1^int^ “stem-like” exhausted T cells that expand during PD-1 pathway blockade. Taken together, our results suggest that Tim-3 is actually more similar to co-stimulatory receptors that are upregulated after T cell activation, rather than a dominant inhibitory protein like PD-1. These findings have significant implications for the development of anti-Tim-3 antibodies as therapeutic agents.

**Significance:** During a chronic viral infection, prolonged exposure to viral antigens leads to dysfunction or “exhaustion” of T cells specific to the virus, a condition also observed in T cells that infiltrate tumors. The exhausted state is associated with expression of specific cell-surface proteins, some of which may inhibit T cell activation. Expression of Tim-3 is associated with acquisition of T cell exhaustion, although it is also expressed transiently during acute infection. Here we provide evidence that a major function of Tim-3 is to enhance T cell activation, during either acute or chronic viral infection. However, Tim-3 is not required for development of exhaustion. Thus, we propose that Tim-3 would be better described as a stimulatory, rather than inhibitory, protein.

---

Tim-3 was first identified as protein specifically expressed by T helper type 1 (Th1) but not T helper type 2 (Th2) T cells (3), but has since been described on multiple of different cell types. More recently, Tim-3 has been most intensively studied for its presence on “exhausted” or dysfunctional T cells, in the presence of persistent antigen (4), including solid and liquid tumors and chronic viral infection (5-7). In addition to Tim-3, other checkpoint receptors such as PD-1, LAG-3, CTLA-4 and TIGIT are highly expressed on exhausted T cells. There are previously described mechanisms by which at least some of these checkpoint receptors may affect T cell exhaustion and/or activation, the intrinsic mechanistic functions of Tim-3 on these cells are largely unknown. Blocking antibodies targeting CTLA-4 and PD-1 are immunotherapy modalities approved for use in humans, but both still only benefit a subset of patients. Therefore, it is critical to understand the function of other checkpoint receptors, like Tim-3, as possible targets of mono- or combination therapy.

During and after acute infection, when antigen is eventually cleared, homeostatically maintained memory T cells are generated. While Tim-3 expression is not well described on memory T cells, it is known that homeostatic cytokines such as IL-2, IL-7 and IL-15 can drive Tim-3 expression (8). However, it is still not clear what possible role Tim-3 may play in the function or development of memory T cells. In a non-human primate model of HIV, Tim-3 is expressed on effector memory T cells, but absent from central memory T cells (9). In addition, it has been shown that during chronic antigen exposure, there is a depletion of memory T cell pools (10). Exhausted T cells are also known to respond poorly to homeostatic cytokines and instead appear to be “addicted” to antigen for survival (reviewed in (11)).

Tim-3 has been identified, either directly or indirectly, as an inhibitory molecule by several studies. However, research into the precise role of Tim-3 on T cells has yielded conflicting data and conclusions. Expression of Tim-3 is correlated with poor viral control and weak recall responses in HIV and HCV infected patients (12, 13). Tim-3 is also present on the most dysfunctional tumor-infiltrating T cells in patients with cancer. Antibodies specific for Tim-3 can result in enhanced T cell function in some settings (14, 15). On the other hand, there is also evidence that Tim-3^+^ T cells are the most responsive cells in active tuberculosis (16). In addition, using germ-line Tim-3 KO mice, a previous study suggested that Tim-3 is necessary for an optimal CD8^+^ T cell response to *Listeria monocytogenes*, although the mechanistic basis for this was not described (17). *In vitro* work from our own lab corroborates a positive role for Tim-3 on T cells where ectopic expression of Tim-3 resulted in increased phosphorylation of multiple positive-acting signaling molecules, and enhancement of downstream T cell activation (18). Finally, studies on other leukocytes, including mast cells and dendritic cells (19-22), are consistent with a potential positive role for Tim-3 in cellular activation.

To help address the discrepancies outlined above, we developed a novel Cre-inducible mouse model that expresses Tim-3 without the need for chronic antigen stimulation. We also employed a recently described Tim-3 KO mouse model (17). Using these tools, we investigated the intrinsic role of Tim-3 in acute and chronic viral infection and found a previously undescribed role for Tim-3 in the formation of effector vs. memory T cells. Our findings suggest a more complex role for Tim-3, based in part on an early co-stimulatory effect.

## Results

Expression of Tim-3 is associated with enhanced TCR signaling and T cell activation. Previous studies have yielded conflicting results regarding the proximal effects of Tim-3 on different cell populations (19, 20, 23, 24). To more precisely assess the relationship between T cell activation and Tim-3 expression, we purified and stimulated naïve T cells from Nur77^GFP^ reporter mice (25). Expression of GFP in leukocytes from these mice is driven by antigen receptor signaling and is proportional to the strength of such signaling (25). As suggested by previous reports (3, 26), Tim-3 expression was detected following *in vitro* stimulation of naïve CD8^+^ T cells. This expression was considerably more robust, and sustained, after at least three rounds of stimulation (Fig. 1A). Furthermore, after two rounds of stimulation, resting CD8^+^ T cells expressing the highest levels of Tim-3^+^ (with or without co-expression of PD-1) had significantly greater expression of Nur77^GFP^. After a third round of stimulation, CD8^+^ T cells uniformly became Tim-3^Hi^ and induced Nur77^GFP^ to a high level (Fig. 1A). Although Tim-3 expression was less robust, a similar trend was observed after two rounds of stimulation (Fig. S1).

**Fig. 1.**
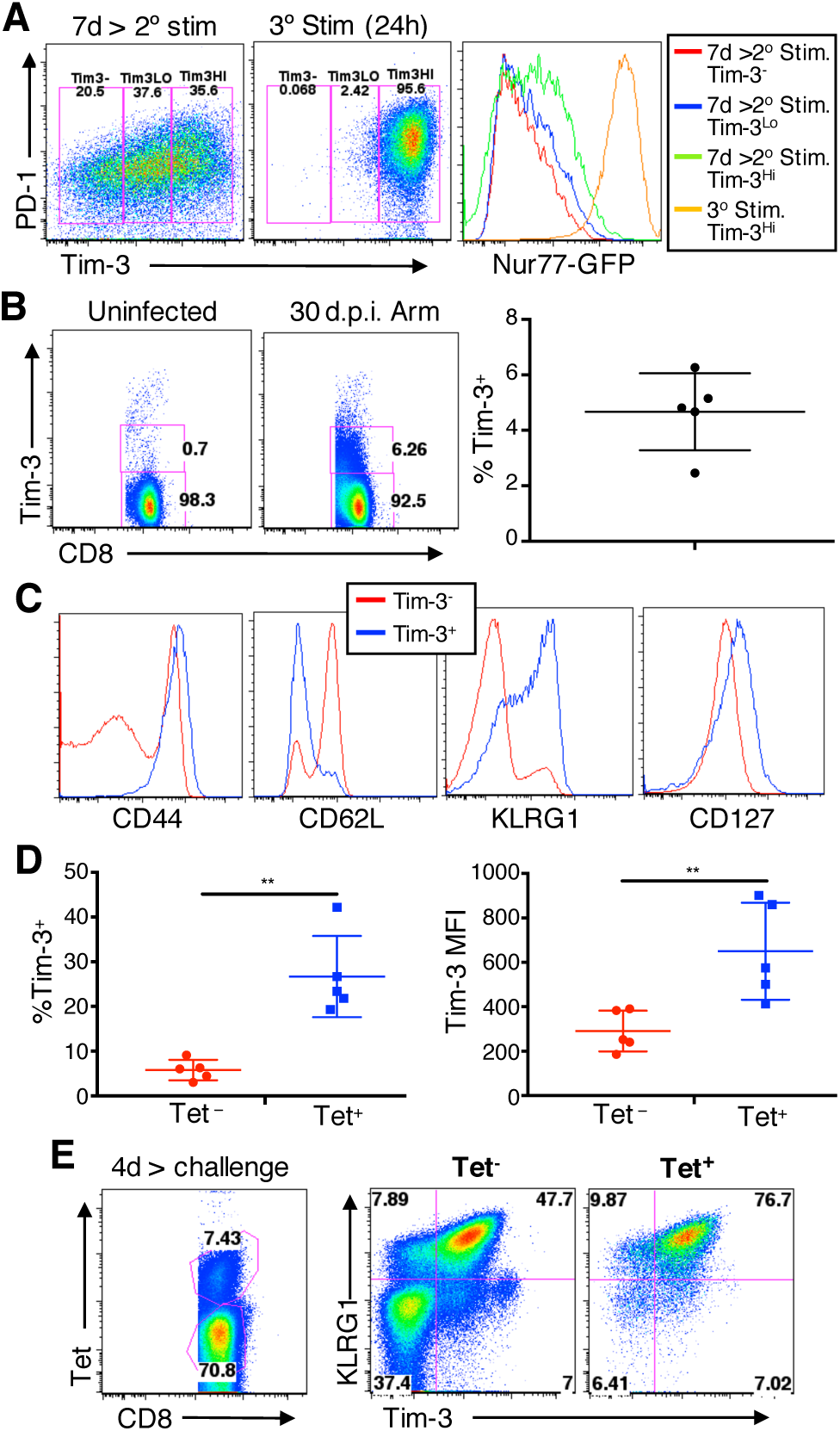
Endogenous Tim-3 expression is associated with increased T cell activation. **A**, CD25-depleted T cells from Nur77^GFP^ mice were stimulated for three days with anti-CD3/CD28 mAbs, followed by a seven day rest with IL-2. After three rounds of stimulation, cells were stained for PD-1 and Tim-3 and analyzed by flow cytometry. Representative of 5 technical replicates per experiment, repeated with n=5 mice. **B**, WT C57BL/6 mice were infected with LCMV Arm. After 30d, spleens were harvested for flow cytometry (n=5 mice, mean ± SD). representative of three independent experiments **C**, Tim-3^+^ vs. Tim-3^−^ CD8^+^ cells were further analyzed for expression of activation and differentiation markers shown in the histograms. **D**, splenocytes from the same experiments as panels b-c were stained with LCMV tetramers plus *α*Tim-3. (n=5 mice, mean ± SD) representative of three independent experiments. **p<0.01 by two-tailed paired Student’s *t* test. **E**, C57Bl/6 mice previously infected with LCMV-Arm (>30d.p.i.) were challenged with LM-GP33 and Tet^−^ vs. Tet^+^ CD8^+^ splenocytes were analyzed for KLRG1 and Tim-3 four days post-challenge. Representative of three independent experiments (n=5).

Expression of Tim-3 and other co-expressed “checkpoint” receptors are usually associated with chronic T cell activation in settings of persistent viral infection or tumors (12-15). We examined the phenotype of T cells that express Tim-3 after acute viral infection, using LCMV Armstrong (Arm) (27). Previously, it was shown that infection of C57BL/6 mice with LCMV Armstrong led to an increase in the percentage of Tim-3^+^CD8^+^ T cells during the acute phase (14). However, we found that a small percentage of Tim-3^+^ cells remain long after the infection is cleared (Fig. 1B). These Tim-3^+^ T cells expressed uniformly high levels of CD44 and KLRG1, a moderately high level of CD127 (IL-7R*α*) and a low level of CD62L (Fig. 1C). This phenotype is consistent with T cells that are antigen-experienced and support the idea that Tim-3 is a marker of activated T cells, although levels of Tim-3 may vary, depending on degree of activation. At day 30 post-infection, we found an increase in both the percentage of T cells expressing Tim-3^+^, as well as its level of expression on tetramer^+^ cells, compared to the tetramer^-^ population (Fig. 1D). Furthermore, upon challenge with *Listeria monocytogenes* GP33 (LM-GP33), the tetramer^+^CD8^+^ cells were uniformly Tim3^+^KLRG1^+^ (Fig. 1E). These data suggest that Tim-3 may enhance activation of antigen-specific T cells both during acute stimulation and also within the memory compartment.

### Tim-3 deficiency results in impaired acute and recall responses by CD8+ T cell

Given that Tim-3 expression is induced by acute LCMV infection and more generally during T cell activation, we tested whether expression of Tim-3 is required to clear LCMV acute viral infection. We infected WT or Tim-3 KO mice with LCMV-Arm and found that by day eight post-infection, the virus was effectively cleared from both groups of mice, with no detectable virus in the spleen. We next assessed the phenotype of the memory T cells formed after infection of WT or Tim-3 KO mice. At day 30 post-infection, Tim-3 KO mice had significantly fewer CD8^+^CD44^+^ T cells, suggesting that fewer T cells were activated during the initial infection (Fig. 2A; Fig. S2A). Consistent with this, there were also fewer antigen-specific T cells remaining in Tim-3 KO mice, using a pool of three LCMV Class-I tetramers (Fig. 2B; Fig. S2B). In addition, fewer of the antigen-experienced CD8^+^ T cells from Tim-3 KO mice upregulated KLRG1, a marker of short-term effector cells (Fig. 2C). Intriguingly, however, this pool of antigen-experienced cells contained a higher percentage of cells expressing CD127 (Fig. 2C), a marker of memory precursor T cells. Upon re-stimulation with LCMV peptides *ex vivo*, fewer Tim-3 KO CD8^+^ T cells responded, based on the proportion of cells producing IFN*γ* and TNF*α* (Fig. 2D; Fig. S2C). Finally, we challenged mice that had cleared Armstrong with *Listeria monocytogenes* expressing GP33 (LM-GP33), an immuno-dominant epitope of LCMV. During the acute recall response, T cells lacking Tim-3 also responded significantly worse than did WT T cells, at the level of cytokine production, degranulation (CD107a exposure) and signaling through the Akt/mTOR pathway, as determined by pS6 staining (Fig. 2E). Thus, in the absence of Tim-3, fewer T cells respond to antigenic stimulation after primary or secondary infection, possibly due to less efficient activation of the Akt/mTOR pathway.

**Fig. 2.**
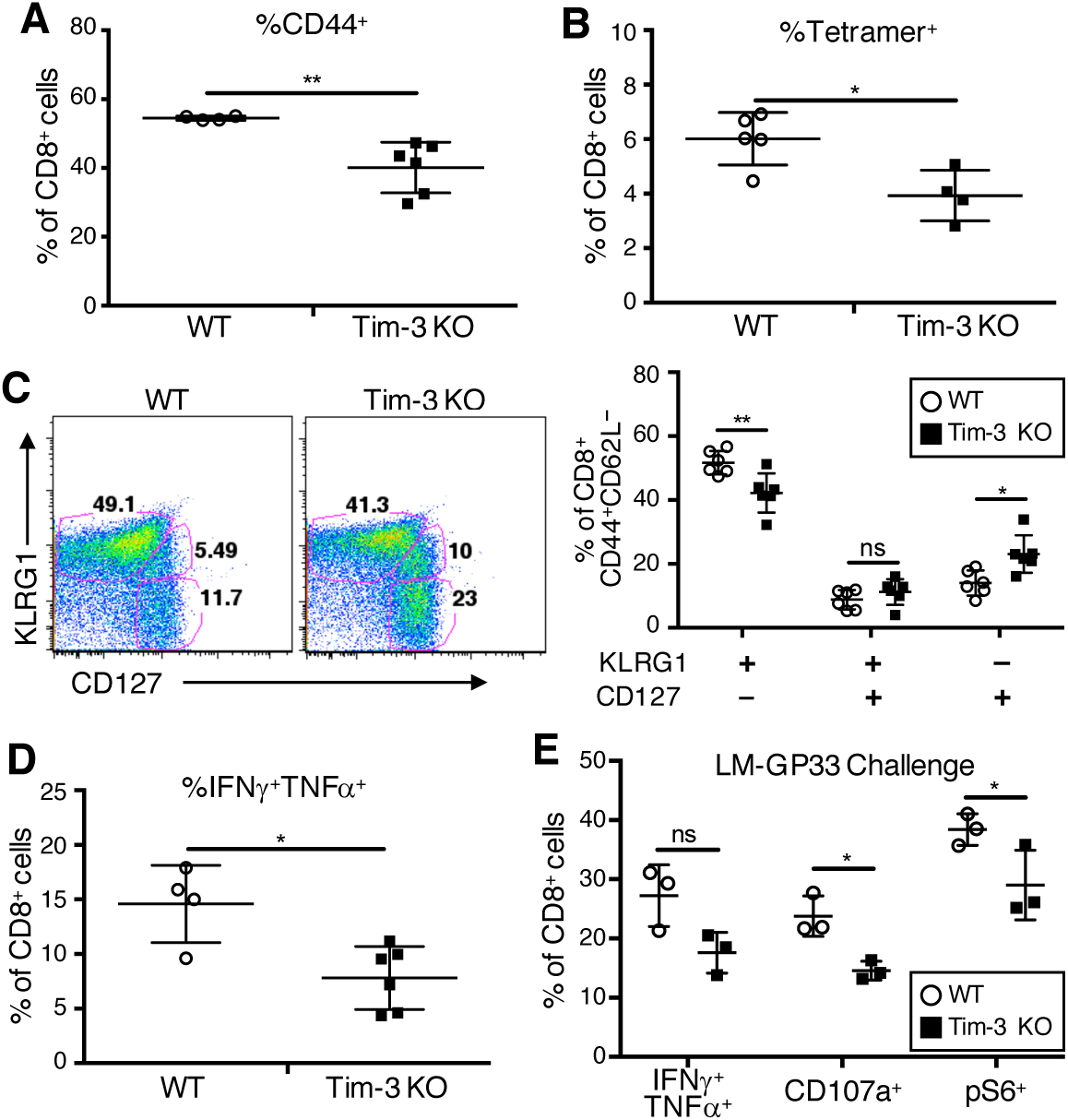
Tim-3 is required for optimal acute responses to primary and secondary infection. Splenocytes from mice infected with 2x10^5^ PFU LCMV-Armstrong were harvested ≥30 d.p.i.. Single cell suspensions were processed and stained for flow cytometry. **A**, %CD44^+^ cells. **B**, Staining for LCMV-specific T cells, using pooled tetramers (GP33, GP276, NP396). **C**, effector vs. memory populations. **D**, Response of splenocytes to stimulation with 100ng/ml pooled LCMV peptides for five hours, in the presence of golgi plug, followed by intracellular cytokine staining and flow cytometry. **E**, cytokine production, degranulation and pS6 staining of *in vitro* stimulated T cells after LM-GP33 challenge. All groups are gated on live CD8^+^ cells. Flow plots are representative of three independent experiments; in the graphs, each point represents an individual mouse with mean ± SD. *p<0.05 **p<0.01 by two-tailed unpaired Student’s *t* test. Representative flow plots for **A**, **B**, and **D** in **Fig. S2.**

### Tim-3 is not required for acquisition of functional exhaustion during chronic infection

Sustained expression of Tim-3 on a subset of responding T cells is associated with the acquisition of T cell exhaustion during chronic infection with LCMV Clone 13. To determine whether Tim-3 expression is necessary for the development of T cell exhaustion, we infected WT and Tim-3 KO mice (17) with Clone 13 virus. Somewhat surprisingly, we observed that Tim-3 KO mice lost significantly more weight and took longer to recover than did WT mice (Fig. 3A). Although there was no significant difference in viral titer in the blood (Fig. 3B), there were significantly fewer antigen-specific T cells in Tim-3 KO mice (Fig. 3C). Chronic infection with LCMV Clone 13 is well-known to induce profound exhaustion among the responding T cells. Somewhat surprisingly, we observed that Tim-3 KO T cells had even fewer CD8^+^ T cells responding by producing IFN*γ* and TNF*α* after peptide re-stimulation (Fig. 3D). It is important to note that regardless of Tim-3 expression, there were no major differences in the expression of other checkpoint receptors on LCMV-specific T cells (Fig. S3). This is the first demonstration that expression of Tim-3 is not required for the development of T cell exhaustion in the LCMV Clone 13 model.

**Fig. 3.**
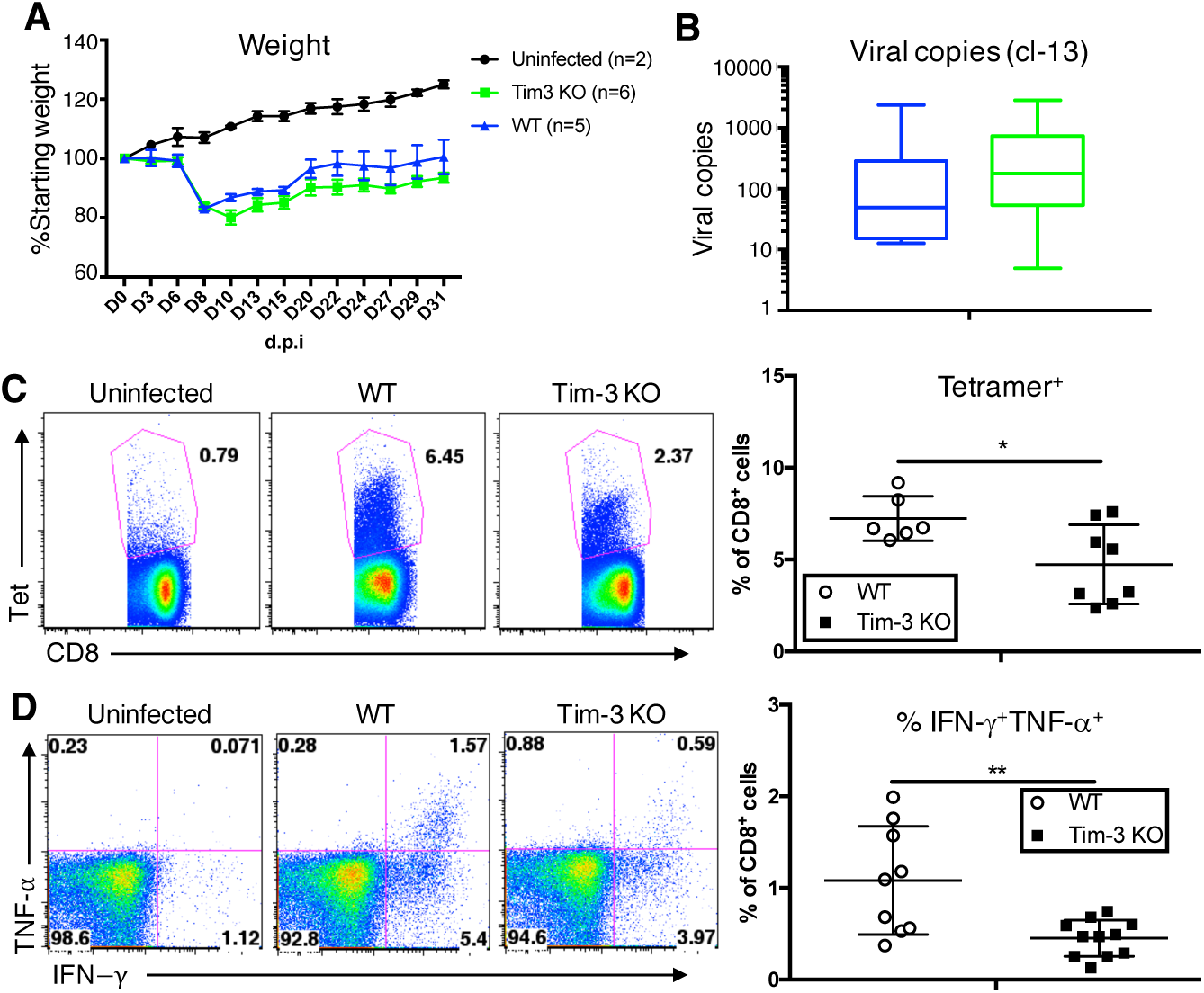
Tim-3 is not required for development of functional T cell exhaustion after chronic viral infection. **A**, Mice were infected with 2x10^5^ PFU LCMV Clone 13 and weighed throughout the course of infection. Uninfected n=2, WT n=5, Tim-3 KO n=6. *p≤0.01 by two-tailed unpaired Student’s *t* tests between WT and KO at that time point (mean ± SD). **B**, At ≥d30 mice from **(A)** were sacrificed and virus in the blood was measured by qPCR using a plasmid standard curve; presented as mean ±SEM. **C**, Tetramer^+^ cells in the spleen were quantified using pooled LCMV tetramers (GP33, GP276, NP396), each point represents an individual mouse, (mean ± SD). Representative of three independent experiments. **D**, Splenocytes were stimulated with 100 ng/ml pooled peptides for five hours in the presence of GolgiPlug, then processed for intracellular staining and flow cytometry analysis, each point represents an individual mouse, (mean ± SD) pooled from three independent experiments. in **C** and **D**, *p<0.05 **p<0.01 by two-tailed unpaired Student’s *t* test.

### Enforced expression of Tim-3 enhances generation of short-term effector T cells

Current models for studying Tim-3 on T cells require persistent antigen stimulation or chronic infection, due to the fact that Tim-3 is generally not stably expressed on the surface of T cells until after several rounds of stimulation (26). A complicating factor in these models is that other checkpoint receptors (e.g. PD-1, TIGIT and LAG3) are also induced with Tim-3. Thus, we developed a flox-stop-flox Tim3 (“FSF-Tim3”) mouse model to drive Tim-3 expression in a Cre-dependent manner (Fig. S4). Upon expression of Cre, the stop cassette is removed, and a flag-tagged version of the murine Tim-3 cDNA is expressed. This model not only circumvents the need for chronic stimulation, but also allows us to enforce Tim-3 expression specifically on T cells to validate our *in vitro* data (18). When FSF-Tim3 mice were bred with CD4-Cre mice, Tim-3 was highly expressed on all *α β* T cells, with no overt effects on T cell development in the thymus or periphery (Fig. S4). This was also the case when FSF-Tim3 mice were bred with E8i-Cre mice, yielding expression of Tim-3 only on CD8^+^ T cells (Fig. S5).

To determine if Tim-3 alone could drive T cell exhaustion during an acute response, we infected FSF-Tim3/CD4-Cre mice, or CD4-Cre control mice, with LCMV-Arm and monitored viral titer in the spleen. FSF-Tim3/CD4-Cre mice cleared LCMV-Arm with the same kinetics as mice expressing CD4-Cre alone (data not shown), however they had significantly more KLRG1^+^ T cells in the antigen-experienced pool (CD8^+^CD44^+^CD62L^−^) at 30 d.p.i. (Fig. 4A). This was accompanied by a lower percentage of KLRG1^+^CD127^+^ T cells, indicating that Tim-3 induction promotes the formation of more short-lived effector T cells at the expense of the long-lived memory pool. Although memory-precursor KLRG1^+^CD127^+^ T cells generally have higher Tbet and lower Eomes expression, KLRG1^+^CD127^+^ memory T cells from enforced Tim-3 expressing cells have an even higher Tbet:Eomes ratio (Fig. 4B). This appears to be the result of decreased Eomes expression in Tim3-expressing cells, as Tbet expression remained unchanged (Fig. 4B). When FSF-Tim3/CD4-Cre and CD4-Cre-only memory T cells were stimulated *ex vivo* with pooled LCMV peptides, there was no significant difference in the percentage of cells producing cytokines (Fig. 4C). However, Tim3-expressing CD8^+^ T cells had greater increase in pS6 staining following peptide stimulation, compared to control T cells, indicating enhanced activation (Fig. 4D).

**Fig. 4.**
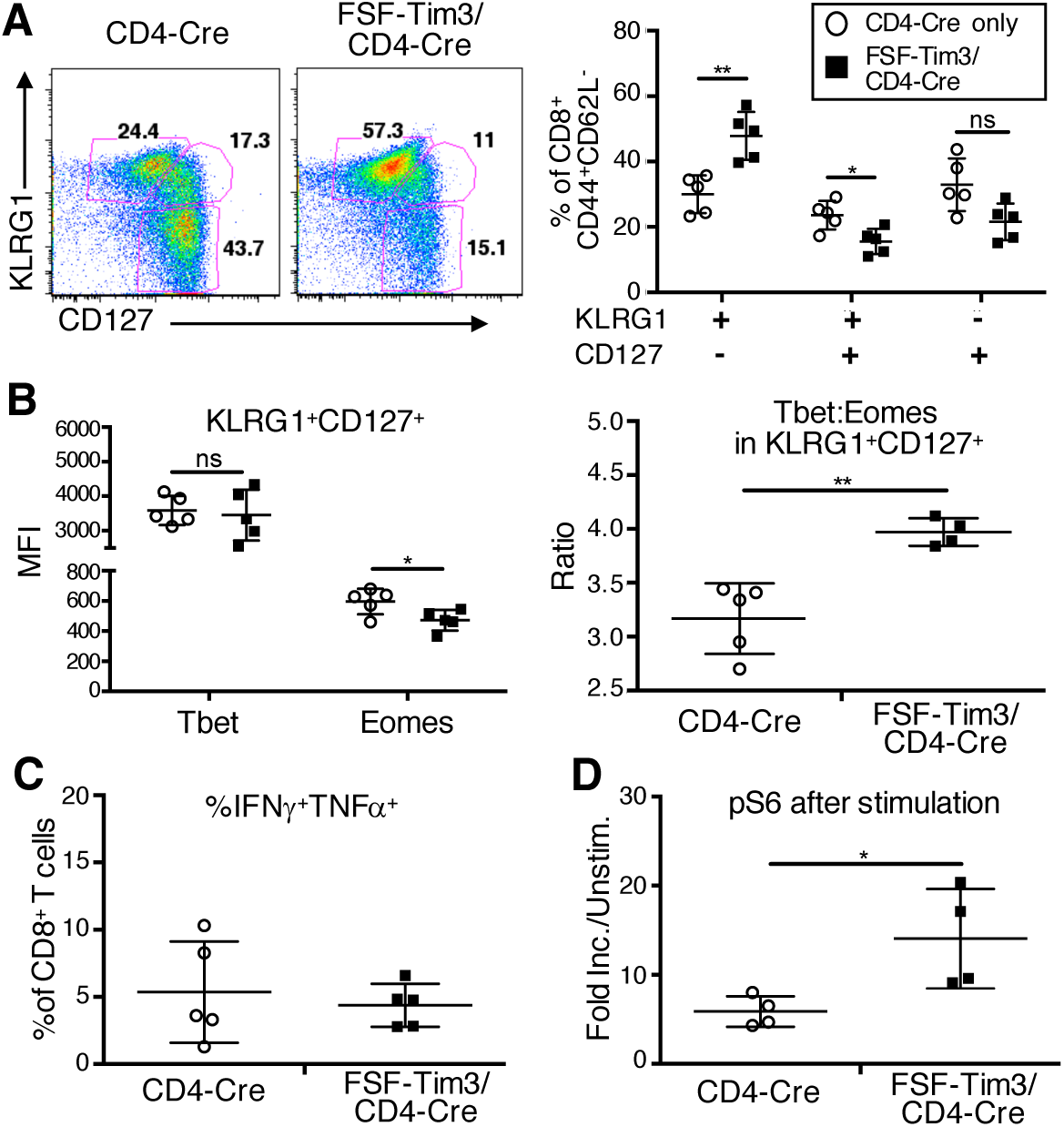
Tim-3 promotes formation of terminal effector phenotype CD8^+^ T cells and enhances early T cell activation. Splenocytes from mice infected with 2x10^5^ PFU LCMV-Armstrong were harvested ≥30 d.p.i. Cells were processed and stained for flow cytometry. **A**, KLRG1 and CD127 were examined in the live antigen-experienced (CD8^+^CD44^+^CD62L^−^) population. **B**, The ratio of Tbet:Eomes (right) was plotted based on MFI of the two (left) in the CD8^+^CD44^+^CD62L^−^KLRG1^+^CD127^+^ population. **C-D**, Splenocytes were stimulated with pooled LCMV peptides for five hours and processed for intracellular staining and flow cytometry of IFN*γ*/TNF*α* (**C**) or pS6 (**D**). Data are representative of three independent experiments; in the graphs each point represents an individual mouse (mean ± SD). *p<0.05 **p<0.01 by two-tailed unpaired Student’s *t* test.

### Tim-3 enhances TCR signaling in a cell-intrinsic fashion

Based on data obtained after LCMV infection (above) and our previously published cell line data (18), we decided to further investigate the effects of Tim-3 on TCR signaling, here using primary T cells from our FSF-Tim3 mice. Intriguingly, the effect of Tim-3 on TCR signaling may occur at a very proximal level, as we observed enhanced induction of phospho-tyrosine after *ex vivo* stimulation of T cells from FSF-Tim3/CD4-Cre mice with anti-CD3/CD28 mAbs (Fig. 5A). We observed even more striking effects when we looked further downstream, at Akt/mTOR signaling. Thus, both pAkt and pS6 were dramatically enhanced in T cells with ectopic Tim-3 expression (Fig. 5B-C). The latter is consistent with the decreased pS6 noted above in LCMV-experienced Tim-3 KO T cells and increased pS6 in FSF-Tim3 mice, also after LCMV infection. To help rule out the possibility of confounding developmental effects of the Tim-3 transgene driven by CD4-Cre, we also investigated activation of CD8+ T cells from FSF-Tim3 mice crossed to E8i-Cre, where expression is specific to CD8 single positive T cells (28). We purified peripheral T cells from these mice and stimulated them in vitro with anti-CD3/CD28, and noted that only the CD8+ T cells in this case responded with enhanced pS6, while CD4+ T cells had equivalent levels of pS6, regardless of whether they came from FSF-Tim3 mice (Fig. 5D). Finally, we purified T cells from WT or FSF-Tim3 mice that did not carry a Cre transgene, and treated these cells overnight with Tat-Cre fusion protein, which induced Tim-3 expression on a subset of T cells from FSF-Tim3 mice (Fig. 5E). The cells were then stimulated overnight with CD3/CD28 mAbs, and analyzed for pS6 and CD69 expression. Consistent with results obtained in the other models, T cells with induced Tim-3 expression responded more robustly to stimulation. Thus, Tim-3 promotes formation of short-term effector T cells, possibly through enhancement of Akt/mTOR signaling and suppression of Eomes expression.

**Fig. 5.**
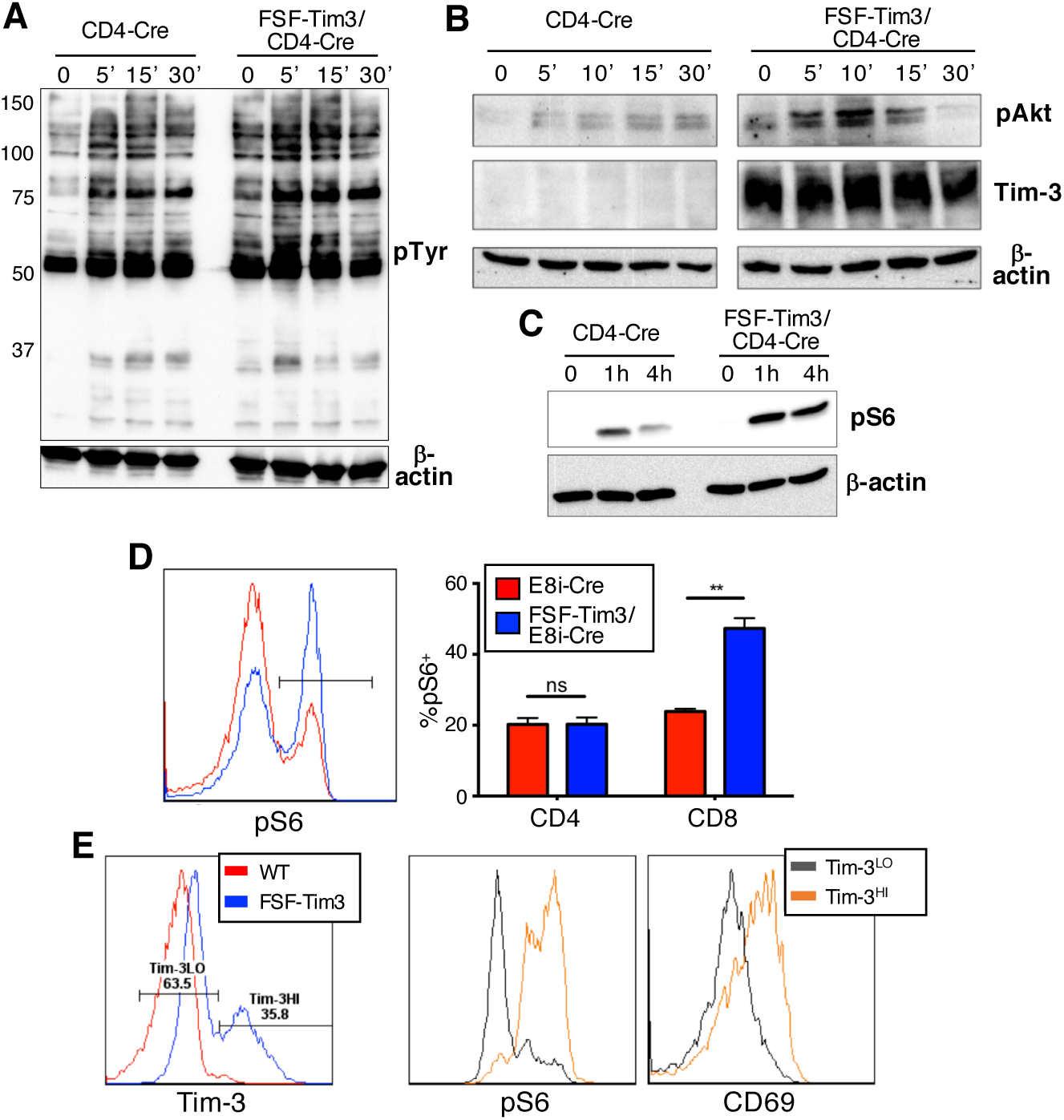
Enhanced TCR signaling in primary T cells with enforced Tim-3 expression. **A**, purified splenic T cells from CD4-Cre or FSF-Tim3/CD4-Cre mice were stimulated *in vitro* with αCD3/CD28 mAb’s for the indicated times. Cells were lysed and analyzed by western blotting for total phospho-tyrosine. Blots were re-probed for β-actin as a loading control (bottom). **B-C**, purified T cells from the indicated mice were stimulated *in vitro* for the indicated times, and analyzed by western blotting for phosphorylation of Akt on S473 (**B**) or for phosphorylation of ribosomal protein S6 (**C**). Blots were also re-probed for Tim-3 (**B**) and β-actin (**B-C**) Blots are representative of three independent experiments. **D**, purified T cells from E8i-Cre or FSF-Tim3/E8i-Cre mice were stimulated *in vitro* for four hours, followed by flow cytometry analysis of pS6. A representative flow cytometry histogram is shown on the left, and compiled data from three experiments is shown on the right (n=3 mice, mean ± SD) **p<0.01 two-tailed unpaired Student’s *t* test. **E,** splenocytes from FSF-Tim3 or a WT littermate were treated with Tat-Cre overnight then stimulated with αCD3/CD28 mAb’s for four hours cells were analyzed by flow cytometry for Tim-3 expression (gated on CD8^+^ live cells). pS6 and CD69 were analyzed within the Tim-3^LO^ and Tim-3^HI^ groups of the FSF-Tim3 cells. Representative of three independent experiments of n=3 biological replicates.

### Tim-3 expression modulates the response to PD-L1 blockade

Recent studies have suggested that antibodies targeting Tim-3 may be useful for the treatment of chronic viral infection or cancer (14, 29). This is due in part to the apparent compensatory upregulation of Tim-3 noted when PD-1/PD-L1 blockade are employed (24, 30). We wanted to determine whether enforced expression of Tim-3 alone would affect the ability of anti-PDL1 antibody treatment to enhance the response to LCMV Clone 13 (31). As shown in Figure 6A, treatment of control (E8i-Cre only) animals with anti-PDL1 mAb modestly reduced viral burden in Clone 13-infected mice, consistent with a previous report (31). Intriguingly, ectopic expression of Tim-3 on CD8+ cells seemed to result in a higher viral load in both isotype Ab control and anti-PDL1 treated animals, relative to the Cre-only controls, although these differences did not reach statistical significance. Recent reports demonstrated that PD-1 pathway blockade in Clone 13-infected mice results in the expansion of a subset of “stem-like” exhausted T cells that express intermediate levels of PD-1, but which are negative for Tim-3, with a relative decrease in PD-1^hi^ cells (32, 33). When we examined the above groups of mice for these PD-1^hi^ vs. PD-1^int^ cells, we noted that in animals with ectopic expression of Tim-3, there were more PD-1^hi^ cells, in both isotype and anti-PDL1 treated mice (Fig. 6B), relative to Cre-only controls. Although anti-PDL1 treatment still led to some expansion of the PD-1^int^ cells (and decrease of the PD-1^hi^ cells), this effect was weaker in the mice with ectopic Tim-3 expression. These results are also consistent with data on the functional capacity of the CD8+ T cells under these conditions, with overall fewer IFN*γ*/TNF*α* producing cells in mice with ectopic Tim-3 expression (Fig. 6C). As with PD-1 expression, a small advantage was conferred on both WT and Tim-3 mice with anti-PDL1 treatment, although this effect was weaker in the mice with ectopic Tim-3 expression (Fig. 6C). These results suggest that Tim-3 may be sufficient to impair the response of exhausted T cells to PD-1 pathway blockade.

**Fig. 6.**
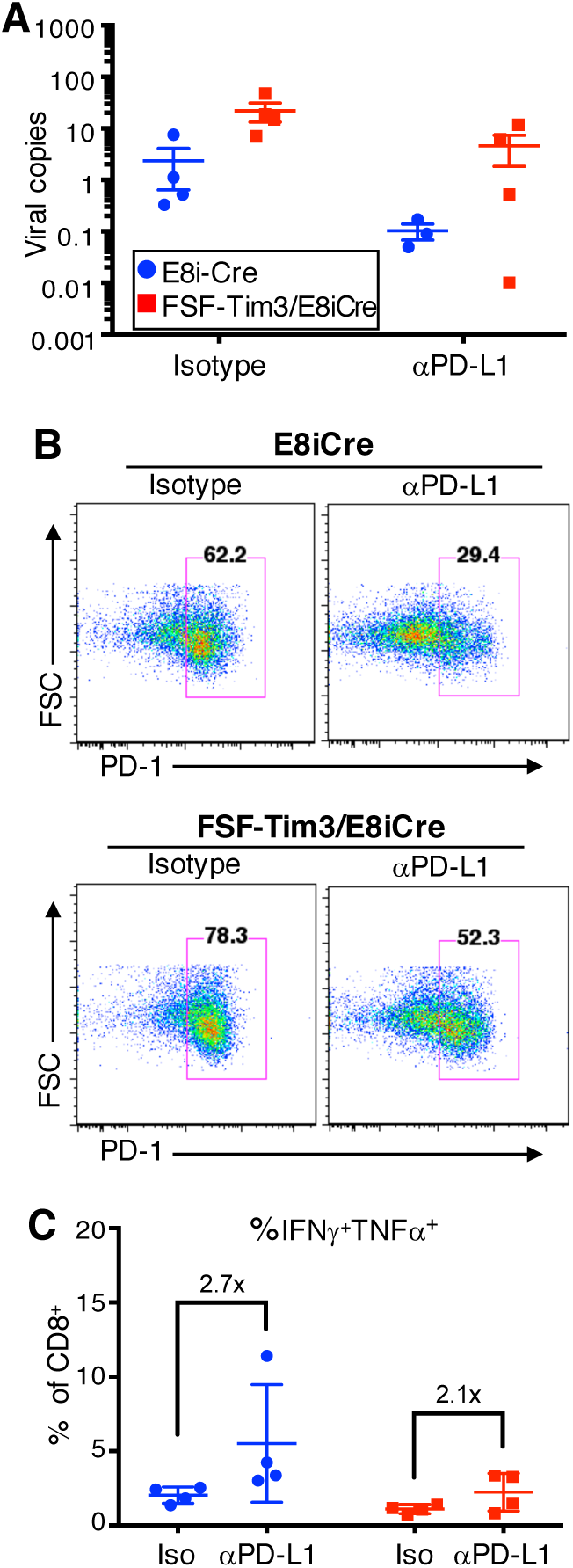
Enforced expression of Tim-3 modulates the response to PD-L1 blockade during chronic LCMV infection. The indicated strains of mice were infected with LCMV Clone 13, and then treated every three days for two weeks beginning 23 d.p.i. with isotype control or anti-PDL1 mAb. Two days after the final treatment (31 d.p.i.) mice were sacrificed and spleens were analyzed. **A**, viral titers in blood 31 d.p.i.. Representative of two independent experiments (mean ± SEM). **B**, Representative flow cytometry analysis of PD-1 expression on splenic CD8^+^ T cells. The gates indicated the location of PD-1^hi^ cells. **C**, quantitation of IFN*γ*^+^TNF*α*^+^ CD8^+^ T cells in spleens of the indicated groups of mice. Each data point represents one mouse (mean ± SD). Representative of two independent experiments.

## Discussion

Although expression of Tim-3 on T cells has been increasingly associated with acquisition of a dysfunctional or exhausted, phenotype, accumulating evidence indicates that Tim-3 is necessary for an optimal effector T cell response. Thus, using genetic models of Tim-3 deficiency or enforced expression, along with infection with LCMV-Arm, we obtained findings consistent with a previous study using *L. monocytogenes* (17). Nonetheless, we believe that ours is the first study to directly link the effects of Tim-3 on early T cell activation and effector T cell development with subsequent impairments in memory T cell formation. The effects of Tim-3 on effector T cell generation during an acute infection correlate well with the early, although transient, expression of Tim-3 after an acute infection (14), and with our own data showing enhanced Nur77^GFP^ activity in Tim-3^+^ cells stimulated *in vitro*. The work reported here suggests a novel mechanism by which Tim-3 could contribute to T cell exhaustion, i.e. by limiting the pool of long-lived memory T cells, while enhancing initial T cell activation and the generation of short-term effector cells (see model in Fig. S6). Depletion of the T cell memory pool is seen in multiple settings where T cell exhaustion has been described, including not just chronic viral infection, but also in the tumor microenvironment (10, 34).

During acute viral infection, expression of Tim-3 and other checkpoint receptors is normally only transient (14). We hypothesized that enforced expression of Tim-3 may cause persistence of an acute infection, although this turned out not to be the case. WT and Tim-3 expressing mice cleared LCMV-Armstrong with similar kinetics, at least at the doses we examined (data not shown). However, enforced expression of Tim-3 did cause an expansion of KLRG1^+^ short-lived effector T cells, at the apparent expense of long-lived CD127^+^ memory precursor T cells. In the transitional population of KLRG1^+^CD127^+^ CD8 T cells, we found that T cells with enforced expression of Tim-3 had a higher Tbet-to-Eomes ratio. This finding provides a possible mechanistic explanation for the expansion of short-lived effector T cells.

In this study, we also found that enforced expression of Tim-3 during acute or chronic LCMV infection had no overt effects on viral pathogenesis, at least at the doses of virus used here. However, the lack of Tim-3 nonetheless resulted in a clear phenotype in chronic LCMV-cl13 infection, as Tim-3 KO mice showed exacerbated weight loss or wasting, which is usually ascribed to immune-mediated pathology (35). Paradoxically, we observed fewer antigen-specific cells, with poor recall response, in these animals. Because this is a germline Tim-3 KO model, it is possible that there is a role for Tim-3 on another cell type or types, which could contribute to the weight loss phenotype. For example, we and others have described functions for Tim-3 in regulating the activation of dendritic cells, mast cells and other myeloid lineage cells (19-22). It should be noted that in this study we did not directly address the function of any of the various reported ligands for Tim-3, including galectin-9, phosphatidylserine (PS), HMGB1 or CEACAM1 (6, 7, 36). An ongoing challenge in this regard is the fact that none of these molecules is an exclusive ligand for Tim-3, making it difficult to ascribe any unique functions to their individual interactions with Tim-3.

Previous studies found that including a Tim-3 Ig with PDL1 further enhances the function of exhausted T cells (14). Therefore, the possibility of combination checkpoint blockade is viable and important to investigate. However, we found it important to first uncover the intrinsic effects of Tim-3 on T cells before determining the effects of ligation. Based in part on studies in which the expression of Tim-3 was associated with reduced effector T cell function in settings of exhaustion, it has been suggested that Tim-3 is a dominant negative regulator of T cell function. Upon closer inspection, however, the effects of Tim-3 on T cell responses appear to be much more complex. Thus, in HIV-infected individuals, Tim-3^+^ T cells have a reduced ability to respond to stimulation, but nonetheless display an enhanced baseline phosphorylation of multiple pathways (37). This is consistent with our finding that murine T cells with enforced Tim-3 expression also have a higher baseline activation, at least based on total phospho-tyrosine levels. In addition, activation-associated phosphorylation events, particularly of pS6 (Ser235/236) and pAkt (Ser473), were enhanced upon polyclonal TCR stimulation above the levels of T cells from Cre-only control animals. Kaech and colleagues found that constitutively active Akt signaling resulted in enhanced KLRG1^+^ expression and ultimately reduced memory T cell formation (38). A similar role for PI3K/Akt signaling in promoting production of short-lived effector cells was reported by Cantrell and colleagues (39). Finally, it has been reported that a downstream target of PI3K/Akt signaling, mTOR, might act to restrict the formation of memory precursor T cells (40).

The work reported here reveals a novel mechanism by which Tim-3 could contribute to T cell exhaustion, i.e. by limited the pool of long-lived memory T cells, while enhancing initial T cell activation and the generation of short-term effector cells. Depletion of the T cell memory pool is seen in multiple settings where T cell exhaustion has been described, including not just during chronic viral infection, but also in the tumor microenvironment. In addition, Tim-3 is necessary for optimal T cell responses in the setting of chronic viral infection. Understanding how Tim-3 contributes to T cell dysfunction in these scenarios will help to guide the development of novel therapies to improve T cell function under what are otherwise suboptimal conditions.

## Materials and Methods

### Mice

Mice were bred in-house under SPF conditions and used between six and eight weeks old. Both male and female mice were used, in equal numbers. For FSF-Tim3 mice, a murine Flag-tagged Tim-3 cDNA that was previously generated in the lab (18) was used to generate knock-in mice. Gene targeting of the *Rosa26* locus with a Flag-Tim-3 construct in C57BL/6 embryonic stem (ES) cells was performed at GenOWay (France). Targeted ES cells were injected, and targeted mice were derived, at U.C. Davis through the MMRC. Mice with a germline KO of Tim-3 were previously described (17), and have been back-crossed to C57BL/6 mice for over 15 generations. CD4-Cre mice on a C57BL/6 background were obtained from Taconic. WT C57BL/6J mice were either obtained from Jackson or bred in-house. All animal procedures were conducted in accordance with the National Institutes of Health and the University of Pittsburgh Institutional Animal Care and Use Committee guidelines.

### Infections and anti-PDL1 treatment

LCMV Clone 13 and Armstrong strains were from Dr. Rafi Ahmed (Emory) and propagated and titered as described previously (41). For acute LCMV-Arm infections, mice were infected with 2x10^5^ PFU intraperitoneally (i.p.). For chronic LCMV-cl13 infections, mice were infected with 2x10^5^ PFU intravenously (i.v.). Infections occurred on day 0 and necropsy on day ≥30 p.i. in age and sex-matched groups. Viral titer was measured from infected tissue by RNA extraction using Trizol LS reagent (Ambion), followed by RT-PCR (Applied Biosystems) and finally qPCR (Applied biosystems) for the GP protein. CT values were compared to a plasmid standard curve, as previously described, to calculate viral copies (42). *Listeria monocytogenes-*GP33 (LM-GP33) was from Dr. Susan Kaech (Yale) and propagated using brain-heart infusion broth as previously described(43). Mice challenged with LM-GP33 received a dose (i.v.) of 2x10^6^ CFU. PDL1 (clone 10F.9G2, BioXcell) or isotype (clone LTF-2, BioXcell) treatment began at d23 after LCMV-Cl13 infection. Mice were randomized to treatment or isotype group and given 200 μg of Ab i.p. every three days for two weeks and harvested two days after the last treatment. The researcher was not blinded to which group the mouse was in.

### Antibodies and Flow Cytometry

Tonbo antibodies: Ghost Dye (Live/Dead) in v510, CD28 (purified) clone 37.51, CD44 (IM7) in APC-Cy7, CD62L (MEL-14) in APC, CD127 (A7R34) in PE-Cy7, CD8 (53-6.7) in FITC or v450, CD4 (GK1.5) in v450, TCRβ (H57-597) in PerCP-Cy5.5, IFN *γ* (XMG1.2) in PE-Cy7. eBioscience antibodies: Eomes (Dan11mag) in PE, CD69 (H1.2F3) in APC, LAG3 (C9B7W) in PE-Cy7, TIGIT (GIGD7) in PerCP-Cy5.5. BD antibodies: TNFα(MP6-XT22) in PerCP-Cy5.5, CD107a (1D4B) in BUV395. Biolegend antibodies: PD1 (RMP1-30) in APC or PE-Cy7, Tbet (4B10) in PerCP-Cy5.5. Cell Signaling Technology: pS6 (Ser235/236) (D57.2.2E) in Alexa 647 or purified. R&D: Tim-3 (215008) in PE, APC or purified. Invitrogen: pAkt (Ser473) (14-6) purified. Millipore: anti-phospho-tyrosine (4G10) platinum. Sigma: β-actin (AC-15) purified. Tetramer staining from pooled LCMV tetramers (NP396, GP276, GP33). MHC peptide monomers, used to make the tetramers, were a gift from Dr. Ahmed (Emory). Samples analyzed for cytokines were fixed/permeablized with BD cytofix/cytoperm kit. Samples analyzed for transcription factors were fix/permeablized with eBioscience FoxP3 staining kit.

### *In vitro* T cell stimulation and western blotting

Splenocytes from infected mice were stimulated with 100ng/mL of pooled LCMV peptides (NP396, GP276, GP33) (Anaspec) unless otherwise noted. Recall stimulations were for five hours at 37 degrees in complete RPMI (Corning) with bovine growth serum (Hyclone). CD25 depletion was done using a CD25 microbead kit (Miltenyi). Lymph nodes from naïve mice were processed from indicated mouse genotype and stimulated with biotinylated CD3 (Tonbo Biosciences) and biotinylated CD28 (eBioscience), cross-linked with streptavidin (Calbiochem) at a 1:1:5 ratio where indicated.

CD3/CD28 stimulation occurred in serum-free RPMI media. Cells were then lysed with RIPA buffer in the presence of protease and phosphatase inhibitors. An SDS-PAGE gel was run and blotted onto PVDF membrane (Millipore), which was blocked with BSA and probed for the indicated proteins.

### Statistical analysis

Samples sizes for *in vivo* experiments were determined based on pilot experiments to be 3-5 mice per group due to the robust phenotype. For *in vitro* experiments, at least four biological replicates were done to ensure statistical power. Variation within and between experiments was within two SD. No data points (or mice) were excluded from analysis unless upon tissue collection there were no viable cells for analysis. Statistics were analyzed using GraphPad Prism software. Paired and un-paired student’s *t* test, and one-way ANOVA were used for data analysis and determination of p-values, as appropriate.

## ACKNOWLDEGEMENTS

We thank R. Ahmed (Emory) for providing viral stocks and protocols; S. Kaech (Yale) for providing LM-GP33 stocks; Katherine Davioli (Univ. Pittsburgh) for propagating LCMV virus; Juan C. de la Torre (Scripps) for the LCMV plasmids used to generate standard curve for viral titer analysis. We are also grateful to T. Hand (Univ. Pittsburgh) and members of the Kane lab for critical feedback and discussion, as well as comments on the manuscript. This work was supported by NIH awards AI109605 and CA206517 to L.P.K., an American Association of Immunologists Careers in Immunology Fellowship, to L.A. and L.P.K., and EY08098 to support Katherine Davioli in the virology & histology core.

## Author Contributions

The project was initially conceived by L.P.K. Experiments were performed by L.A. and A.W. Data were analyzed by L.A. and A.W. The manuscript was written by L.A., with contributions from A.W. and L.P.K.

